# The PCI domains are “winged” HEAT domains

**DOI:** 10.1101/2022.05.05.490838

**Authors:** Eleanor Elise Paul, Assen Marintchev

**Affiliations:** Department of Physiology & Biophysics, Boston University School of Medicine, 700 Albany St. W336, Boston, MA 02118, USA

**Author notes:** Corresponding Author, (AM).

## Abstract

The HEAT domains are a family of helical hairpin repeat domains, composed of four or more hairpins. HEAT is derived from the names of four family members: **<u>h</u>**untingtin, eukaryotic translation **<u>e</u>**longation factor 3 (eEF3), protein phosphatase 2 regulatory **<u>A</u>** subunit (PP2A), and mechanistic **<u>t</u>**arget of rapamycin (mTOR). HEAT domain-containing proteins play roles in a wide range of cellular processes, such as protein synthesis, nuclear transport and metabolism, and cell signaling. The PCI domains are a related group of helical hairpin domains, with a “winged-helix” (WH) subdomain at their C-terminus, which is responsible for multi-subunit complex formation with other PCI domains. The name is derived from the complexes, where these domains are found: the **<u>P</u>**roteasome “lid” regulatory subcomplex, the **<u>C</u>**OP9 signalosome (CSN), and eukaryotic translation **<u>i</u>**nitiation factor 3 (eIF3). We noted that in structure homology searches using HEAT domains, sometimes PCI domains appeared in the search results ahead of other HEAT domains, which indicated that the PCI domains could be members of the HEAT domain family, and not a related but separate group. Here, we report extensive structure homology analysis of HEAT and PCI domains, both within and between the two groups of proteins. We present evidence that the PCI domains as a group have greater structural homology with individual groups of HEAT domains than some of the HEAT domain groups have among each other. Therefore, our results indicate that the PCI domains have evolved from a HEAT domain that acquired a WH subdomain. The WH subdomain in turn mediated self-association into a multi-subunit complex, which eventually evolved into the common ancestor of the Proteasome lid/CSN/eIF3.

## Introduction

Helical repeat domains are widespread in eukaryotic genomes, and involved in virtually every major cellular process. These encompass several families, including tetratricopeptide repeat (TPR), ankyrin repeat (ANK), armadillo (ARM), as well as the HEAT repeat domain family (reviewed in [1–4]). The acronym HEAT is derived from the names of several founding members of the family: **<u>h</u>**untingtin, eukaryotic translation **<u>e</u>**longation factor 3 (e**<u>E</u>**F3), protein phosphatase 2 regulatory **<u>A</u>** subunit (PP2**<u>A</u>**), and mechanistic **<u>t</u>**arget of rapamycin (m**<u>T</u>**OR).[5] Of these families, ARM and HEAT repeat domains are only found in eukaryotes, and likely arose more recently during evolution [2]. Another, smaller group of eukaryote-specific helical repeat proteins are the PCI domains. Most PCI domain-containing proteins are subunits of the **<u>P</u>**roteasome “lid” regulatory subcomplex, the **<u>C</u>**OP9 signalosome (**<u>C</u>**SN), or eukaryotic translation **<u>i</u>**nitiation factor 3 (e**<u>I</u>**F3) [6–9].

Unlike most globular domains, helical repeat domains do not have a fixed size, with a distinct beginning and end. Instead, they consist of varying numbers of helical repeats. The larger domains have varying curvatures and tend to form solenoids. This plasticity likely contributed to the fast divergence of helical repeat domains in size, structure, sequence, and function. Sequence analyses have revealed that it is often impossible to detect sequence homology, even among members of the same domain family. Sequence conservation is limited to the interfaces between adjacent helices and tends to be lower than for other domain families. The variable length, irregular shapes, and low/undetectable sequence homology have made the analysis of the evolution of these domain families challenging (reviewed in [1, 3, 5]).

The HEAT repeat domain family was first reported in 1995[5]; these domains consist of a series of helical hairpins, each formed by two antiparallel helices, packing against each other and the two surrounding helical hairpins (**Fig. 1**). The number of hairpins varies from as few as four to over 50 [5, 10–12].

**Figure 1.**
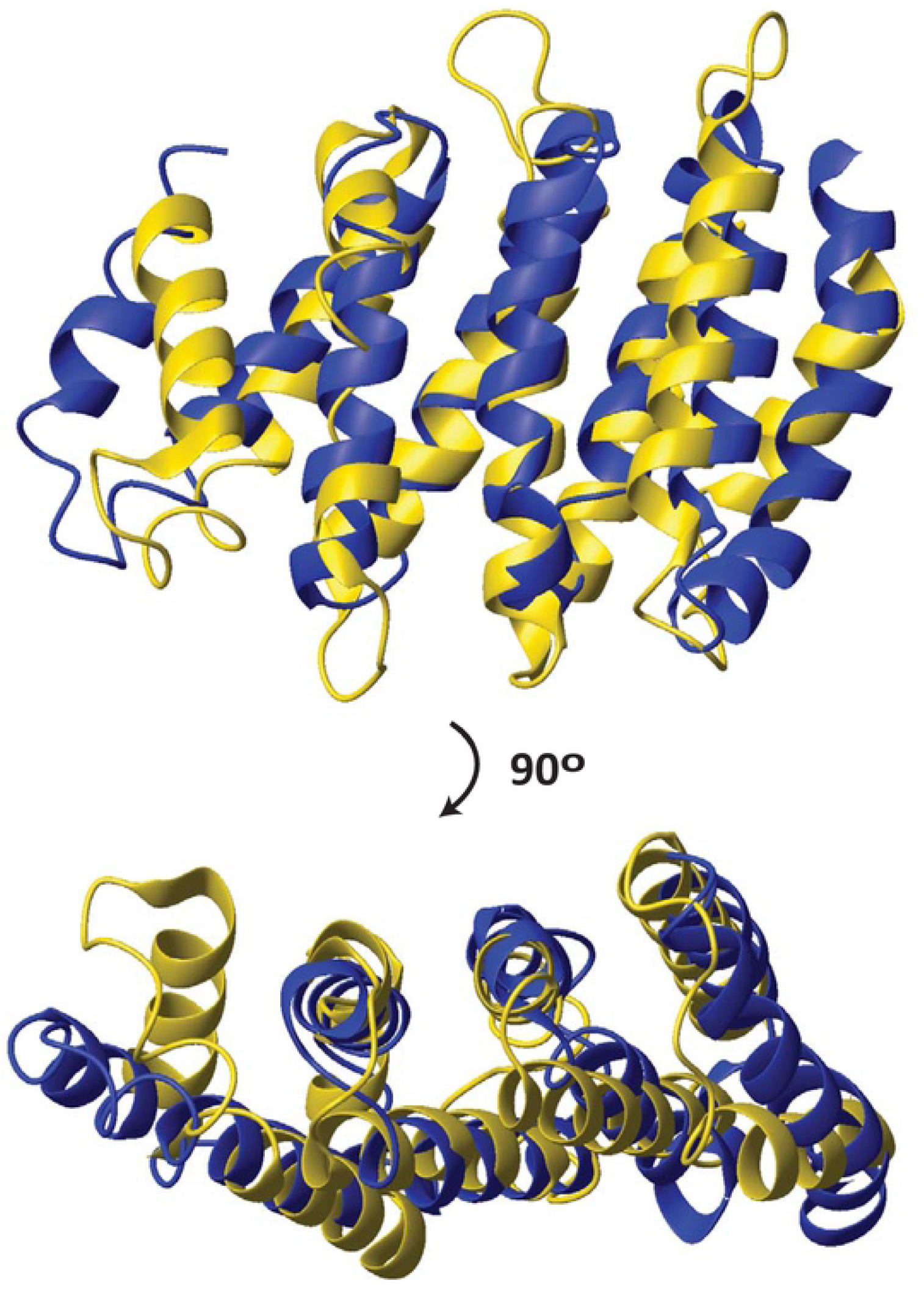
Structure alignment between the HEAT domain of CTF3 and the PCI domain of RPN12. DALI [23]-based structure alignment between the HEAT domain of *S. cerevisiae* CTF3 (6wuc.pdb, chain B) and the PCI domain of human RPN12 (5l4k.pdb, chain P). The structures are shown in ribbon. CTF3 is colored gold; RPN12 is colored blue. Only the aligned portions of the two structures are shown.

PCI domains have very similar organization to the HEAT domains and the other helical repeat domains, but also have a “winged-helix” (WH) subdomain at their C-terminus. The WH subdomains mediate complex assembly with the WH subdomains of other PCI domain containing subunits. The 26S Proteosome lid, CSN, and eIF3 are multiprotein complexes with a common ancestor and similar organization, including the presence in each of them of six PCI domaincontaining subunits: RPN3, RPN5, RPN6, RPN7, RPN9, and RPN12 in the Proteasome lid; CSN3, CSN4, CSN2, CSN1, CSN7, and CSN8 in the CSN; and eIF3l, eIF3a, eIF3c, eIF3e, eIF3m, and eIF3k in eIF3, respectively. The pairwise correspondence of some subunits in each of these complexes was possible to establish based on sequence homology, while for others, it became obvious when the structures of the Proteasome, CSN and eIF3 became available. In each of these complexes, the WH subdomains form a ring in the core of the complex [6–8, 13–15]. eIF3 in some groups of organisms has lost part of its subunits and has fewer than six PCI domains. For example, *Saccharomyces cerevisiae* has two of the PCI subunits: eIF3a and eIF3c; while *Schizosaccharomyces pombe* has four: eIF3a, eIF3c, eIF3e, and eIF3m subunits [16–22].

The structure similarity between HEAT and PCI domains is illustrated in **Fig. 1** and has been noted previously [9], although no sequence homology can be detected. However, as pointed out above, there is no detectable sequence homology even between relatively close homologs within the HEAT and PCI families [8, 14].

When performing DALI [23] searches for structural homologs of HEAT proteins, we noticed that sometimes PCI domain structures appeared in the search results with higher scores that some of the known HEAT domains. This prompted us to investigate whether the PCI domains are a subset of the HEAT family, or the PCI and HEAT domains are two related but distinct families. Here we performed extensive comparative structure homology analysis, which demonstrates that the PCI domains are a sub-family of the HEAT domains, which originated when a HEAT domain acquired a WH domain at its C-terminus.

## Results

We reasoned that if structure homology between the PCI domains and individual groups of known HEAT domains is comparable to the structure homology among the groups of HEAT domains, then the PCI domains are a group of HEAT domains that acquired a WH subdomain during evolution, after the HEAT domains had diverged from other helical repeat domain families. Conversely, if all groups of HEAT domains have greater structure homology to each other than to the PCI domains, then the PCI domains split earlier, before the common ancestor of all HEAT domains (**Fig. 2**):

- If the PCI domains have lower structure homology scores with all groups of HEAT domains than the scores among HEAT domain groups, then the PCI domains split off from their common ancestor with HEAT domains before the individual HEAT domain groups diverged from each other (**Fig. 2A**).
- If the average homology scores between PCI domains and at least one group of HEAT domains are comparable, or higher, than the lowest average homology scores between at least two groups of HEAT domains, then the PCI domains are a group of HEAT domains that have acquired a winged helix (**Fig. 2B**).

**Figure 2.**
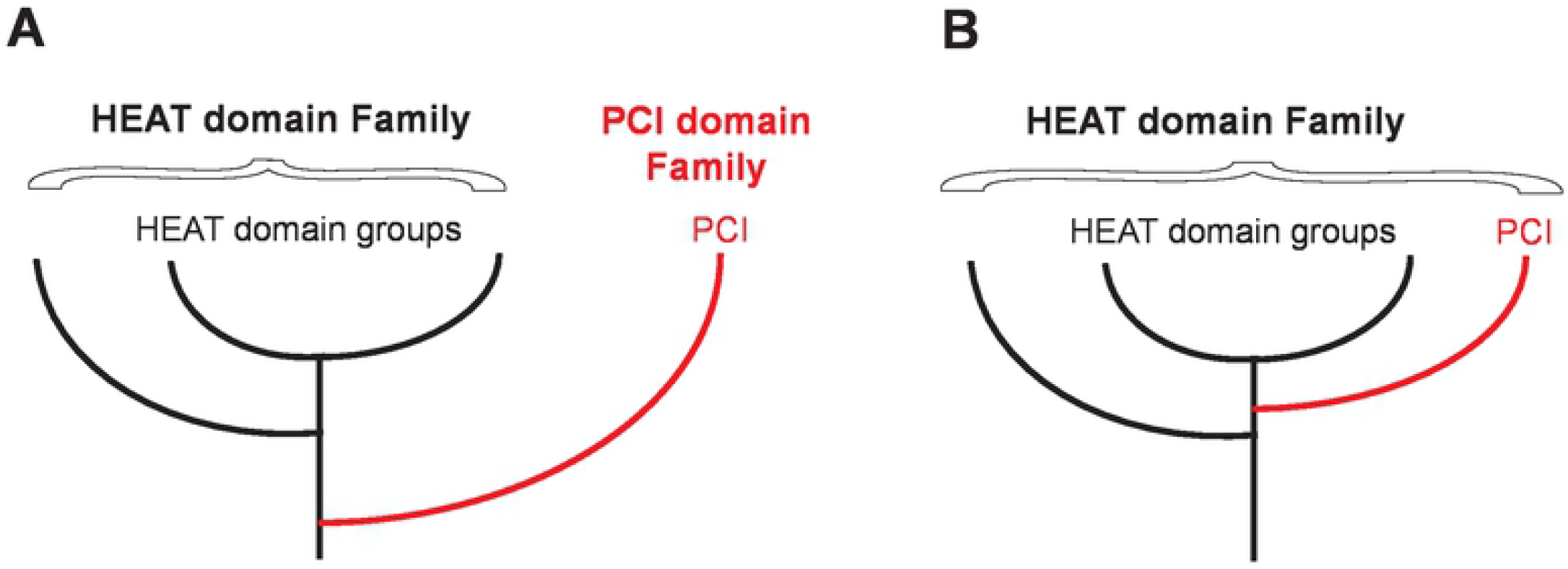
Possible alternatives for the evolution of PCI domains. **A.** PCI domains (red) diverged from HEAT domains before the last common ancestor of all HEAT domains and constitute a separate family of domains, as currently thought. **B.** The PCI domains are members of the HEAT domain family that acquired a WH subdomain after the last common HEAD domain ancestor.

To account for the presence of a Winged Helix (WH) subdomain in PCI domains, but not in the HEAT domains, which could affect the structure homology scores, we also used PCI domain structures with deleted WH subdomains. Finally, helical repeat domains pose unique challenges for structure homology software, because they have a repetitive structure, a variable number of repeats, and varying curvature. To try to compensate partially for these factors, we also used four-hairpin fragments of HEAT and PCI domains.

We performed extensive searches for HEAT and PCI domains with known structures and selected a representative set of structures with high-resolution and avoiding close relatives. We then subdivided the representative structures into groups based on structure homology scores. The HEAT domains fell into four groups, based on structure homology, as well as a couple of outlier structures, which were considered as a fifth group. (**Supplementary Table S1**) The majority of the “founding” members of the HEAT domain family, including Huntingtin, eEF3, and PP2A [5], were in Group 1. The MA3, MIF4G, and W2 domains [10–12, 24–27] formed individual groups, Groups 2, 3, and 4, respectively. Finally, two of the selected proteins, including mTOR, one of the founding HEAT domain family members [5], were outliers and placed in Group 5. The PCI domains formed Group 6.

formed Group 6.

We found that the average homology score between Group 1 HEAT domains and PCI domains (Group 6), was 4.8 (where scores above 2 indicate homology). This score was higher than those between Group 1 and Groups 2 (4.2) and 4 (3.6). Similar results were observed for the outlier HEAT domains (Group 5): average homology score with the PCI domains (Group 6) was 4.6, whereas those with Groups 2 and 4 were both 3.9. The homology score of Group 5 with Group 3 was 4.5, comparable to that with Group 6 (PCI) (**Table 1,** The numbers in Fig. 3 plus intragroup Z-scores **Fig. 3A**). This observation indicates that the PCI domains belong to the family of HEAT domains.

**Table 1.**
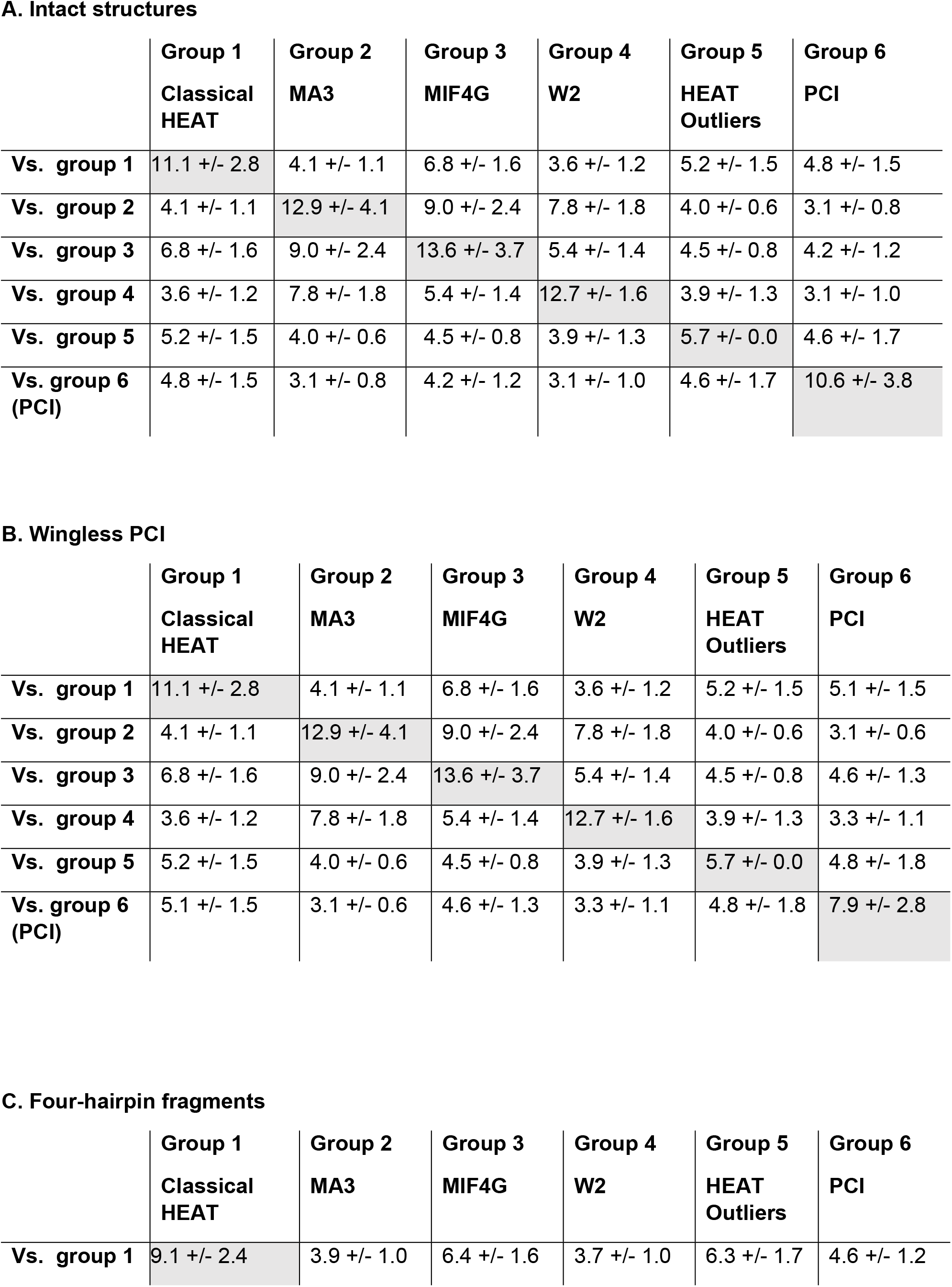
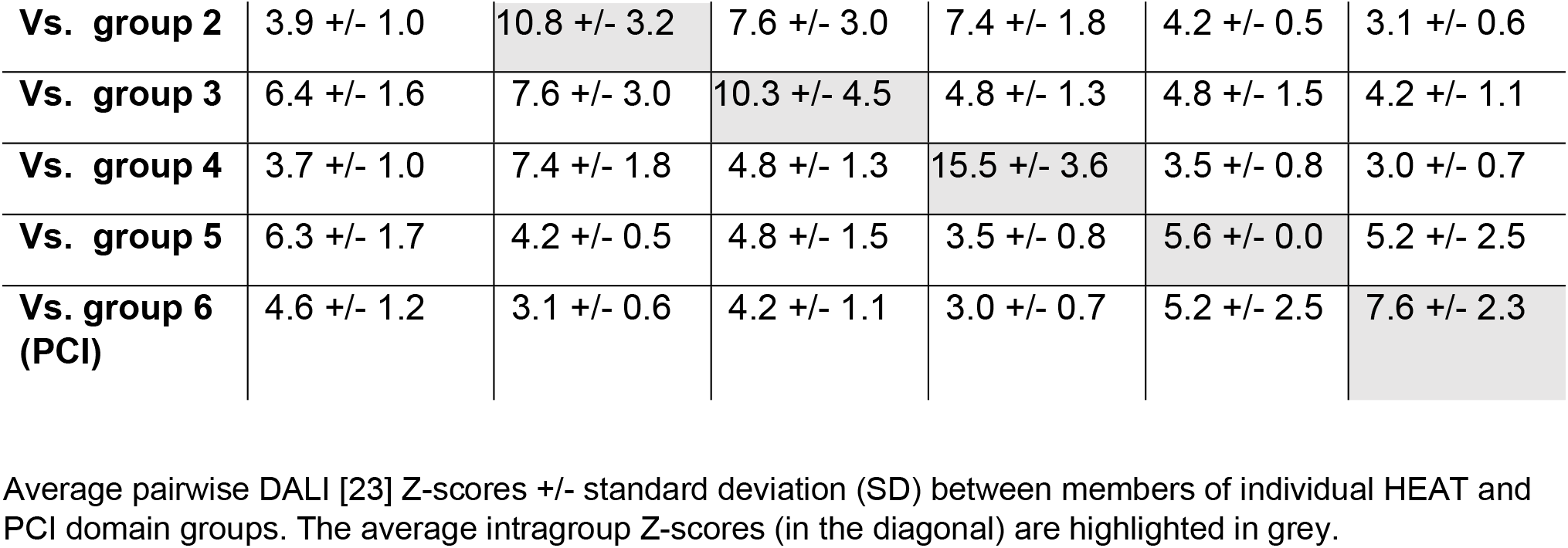
Structure homology scores among groups of HEAT and PCI domains.

**Figure 3.**
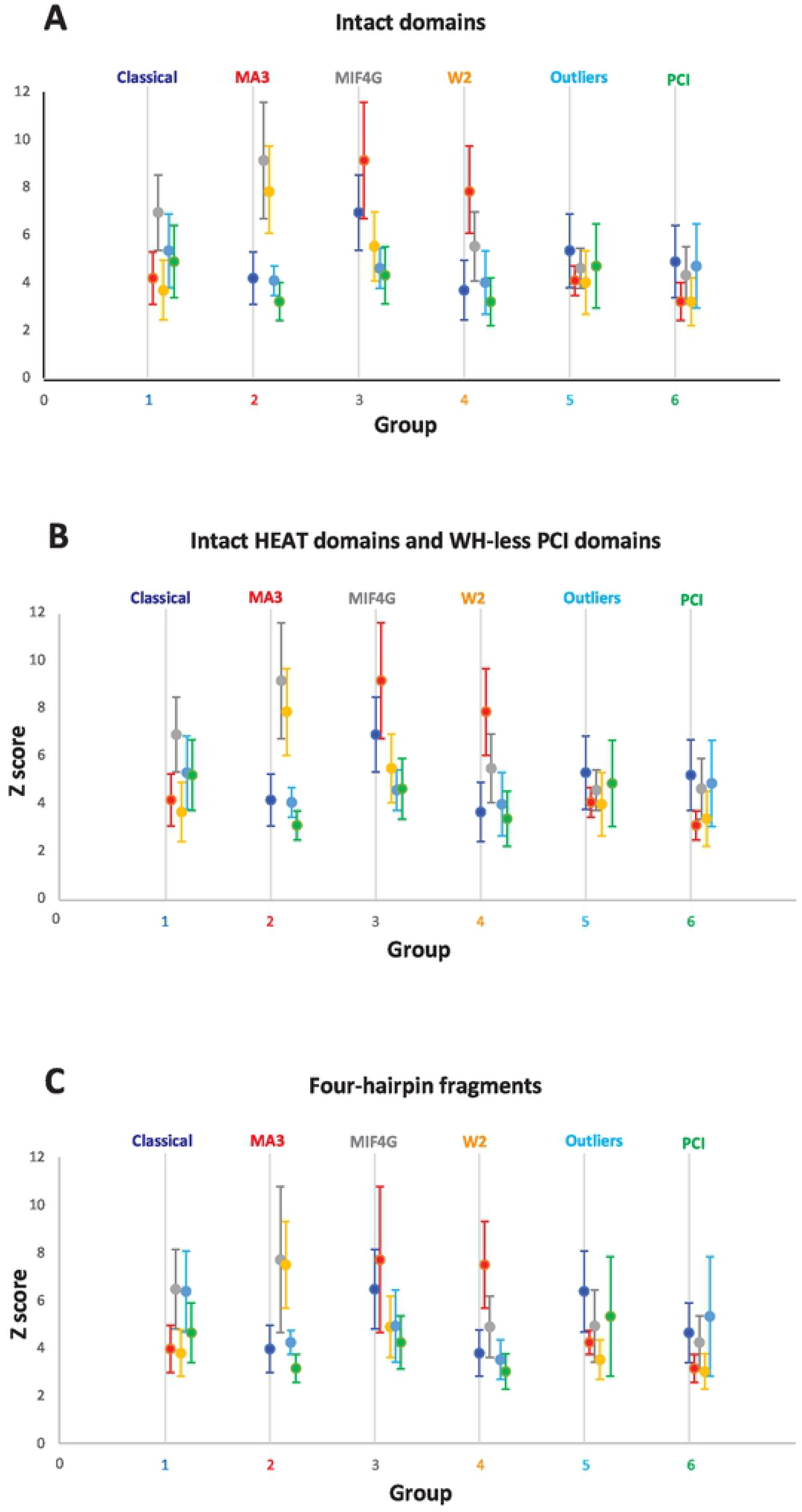
Structure homology scores among groups of HEAT and PCI domains. Average pairwise DALI [23] Z-scores +/− standard deviation (SD) between members of individual HEAT and PCI domain groups. Groups are labeled and color-coded. The Z-scores of each group with the other five groups are shown on the Y-axis, with slight offset along the X-axis, and color-coded. Classic HEAT domains (Group 1) are navy. MA3 domains (Group 2) are red. MIF4G domains (Group 3) are grey. W2 domains (Group 4) are orange. Outlier HEAT domains (Group 5) are light blue. PCI domains are green. Intragroup Z-scores are not shown.

The same conclusion was reached when using a set of PCI domains with deleted WH subdomain, which tended to increase further the homology scores between the PCI domains (Group 6) and the HEAT domain groups. The score between Group 1 and Group 6 was now 5.1, and that between Group 5 and Group 6 was now 4.8. Furthermore, the new score between Group 3 and Group 6 was now 4.6, comparable to that between Group 3 and Group 5 (4.5) (**Table 1, Fig. 3B**).

Finally, we used a set of protein fragments with only four helical hairpins, so that all structures had the same size. Again, the average homology score between Group 1 HEAT domains and PCI domains (Group 6), 4.6, was higher than those between Group 1 and Groups 2 (3.9) and 4 (3.7). Likewise, the average homology score of Group 5 with the PCI domains (Group 6), 5.2, was higher than those with Group 2 (4.2), Group 3 (4.8), and Group 4 (3.5) (**Table 1, Fig. 3C**). Thus, all three versions of the analysis yield the same conclusion: that the PCI domains are a group of HEAT domains that have acquired a WH subdomain (the scenario shown schematically in **Fig. 1B**, above).

## Discussion

The comparative structure homology analysis of HEAT and PCI domain structures presented here shows that the PCI domains are members of the HEAT domain family. In fact, they appear more closely related to the canonical HEAT domains than do the MA3, and W2 domains, while the canonical HEAT domains have similar structure homology with the MIF4G and PCI domains (**Fig. 3**). Sometimes described as atypical HEAT domains [10] and found mostly in eukaryotic translation initiation factors and proteins involved in translation regulation and ribosome biogenesis, the MA3 and W2 HEAT domains tend to be shorter, some with as few as four helical hairpins, often arranged in tandem, and likely evolved from a common ancestor containing a MIF4G, an MA3, and a W2 domains in a row, as observed in eIF4G and CBP80 (reviewed in [28, 29]).

To avoid possible score bias due to the length of individual domains, we repeated the structure homology analysis using four-hairpin fragments of the proteins, which corresponds to the size of the smallest HEAT domains. The results with four-hairpin fragments confirm the results obtained with the intact domains. Using the four-hairpin fragments has its own caveats, because, while for the PCI domains, using the last four hairpins before the WH subdomain ensures that corresponding fragments are used, this is not guaranteed to be the case for the various HEAT domains, even if using the last four hairpins of each protein. However, the observation that the evolutionary trees and groups obtained using intact domains and four-hairpin fragments are very similar, supports the validity of this approach. Importantly, both the analysis using intact domains, and that using four-hairpin fragments yield the same results, which strongly supports the overall conclusions.

The MIF4G and MA3 domain groups are evolutionarily closer to each other than to the rest of the HEAT domains, while the W2 domains are closer to the MA3 domains that to the rest of HEAT domains, including the MIF4G group (**Fig. 3**). These observations indicate that the MA3 domains probably resulted from the duplication of a MIF4G domain, and that the W2 domains, in turn, resulted from the duplication of an MA3 domain, as opposed to breaking up of a long HEAT domain solenoid into three consecutive shorter HEAT domains. The propensity of these short HEAT domains to duplicate is illustrated by UPF2, which has three tandem MIF4G domains, and Pdcd4, which has two MA3 domains [30]. eIF5, eIF2Bε, and 5MP/BZW all have W2 domains closely homologous to the W2 domain of eIF4G [25, 29]. 5MP/BZW also has a predicted MA3 domain N-terminal to the W2 domain (ref. [29] and AlphaFold [31, 32]).

In conclusion, this study offers insights into the evolution of the HEAT domain family in eukaryotes and shows that the PCI domains are in fact HEAT domains, which have acquired a WH subdomain at their C-terminus. This further expands the number of proteins containing a HEAT domain, and the already wide range of functions performed by these domains. It would be interesting to trace the origins and divergence of all helical repeat domain protein families as a whole, throughout the evolution of eukaryotes.

## Methods

### Structure homology searches

The DALI Server [23] was used to search the RCSB PDB database [33] for homologous structures and obtain structural homology scores (Z-scores). The goal was to assemble a diverse set of HEAT and PCI domain structures, as well as identify potential new structures belonging to these families. Searches were initiated with several known HEAT domain and PCI domain structures. An initial diverse set of structures was assembled, eliminating structures with high similarity, as well as selecting high-resolution structures. Each structure and the corresponding structure-based sequence alignment were inspected manually, e.g., to confirm whether a putative PCI domain indeed contains the obligate WH subdomain, or that the DALI-generated automatic structure alignment does indeed define a contiguous helical hairpin structure segment. PyMol [34] was used for structure visualization.

### Structure classification

Any non-HEAT or PCI domain portions of the structures were deleted at this stage using PyMol [34]. Structure homology scores were calculated for all pairs of structures in the dataset, followed by a second round of removing highly similar structures, yielding the final dataset (**Table S1**). Where DALI failed to detect homology, we did not use the pair in future analyses, because the structures are in fact homologous. Using the pairs with a Z score 0, or with a Z score 2 (the minimum score considered statistically significant) did not affect the conclusions.

The evolutionary tree automatically generated by the DALI server served as a starting point for grouping the structures in the dataset. The HEAT domains fell into four groups, “classic” HEAT domains, MA3 domains, MIF4G domains, and W2 domains, all of which had previously been defined [5, 10–12, 24–26, 35]. Two HEAT domain structures were not part of any of the groups and were assigned a separate group (Group 5).The PCI domains formed one group (Group 6). The autogenerated dendrogram was correct in most cases, except grouping CTIF3 with Group 3 (MIF4G), instead of Group 1 (“classic” HEAT domains), and grouping the CBP80 MIF4G domain with Group 2 (MA3), instead of Group 3 (MIF4G). In both cases, the Z scores in **Table S1** unambiguously assign these two structures to the correct group.

### Analysis of structure homology and evolutionary relationships

We calculated average structure homology scores (Z scores) and standard deviations for every pair of groups from the pairwise Z scores between all members of the two groups, obtained using the DALI server [23]. The average Z scores between groups of HEAT domains were compared to those between a HEAT domain group and the PCI domains, in order to determine whether or not at least one of the groups of HEAT domains shows greater, or at least similar, homology to the PCI domains than to at least one other group of HEAT domains.

To evaluate the contribution of the WH subdomain in the PCI domains, we repeated the analysis using “wingless” PCI domains. The WH subdomains were deleted from the corresponding structures in PyMol [34]. To account for the fact that different HEAT and PCI domains vary greatly in size, we generated four-hairpin fragments from each structure in the dataset and repeated the analysis with those. MOLMOL [36] was used to create figures of structure alignments between HEAT and PCI domains.

